# Agent-based models help interpret patterns of clinical drug resistance by contextualizing competition between distinct drug failure modes

**DOI:** 10.1101/2022.02.25.481999

**Authors:** Scott M Leighow, Benjamin Landry, Michael J. Lee, Shelly R. Peyton, Justin R. Pritchard

## Abstract

**Introduction:** Modern targeted cancer therapies are carefully crafted small molecules. These exquisite technologies exhibit an astonishing diversity of failure modes (drug resistance mechanisms) in the clinic. This diversity is surprising because back of the envelope calculations and classic modeling results in evolutionary dynamics suggest that the diversity in the modes of clinical drug resistance should be considerably smaller than what is observed. These same calculations suggest that known microenvironmental resistance mechanisms should not be able to compete for outgrowth with genetic resistance within a tumor, and yet evidence of microenvironmental resistance is often observed in the clinic. Quantitatively understanding the underlying biological mechanisms of failure mode diversity may improve the next generation of targeted anticancer therapies. It also provides insights into how intratumoral heterogeneity might shape interpatient diversity during clinical relapse.

**Materials and Methods:** We employed spatial agent-based models to explore regimes where spatial constraints enable microenvironmental resistance to significantly compete with genetically resistant subclones. In order to parameterize a model of microenvironmental resistance, BT20 cells were cultured in the presence and absence of fibroblasts from 16 different tissues. The degree of resistance conferred by cancer associated fibroblasts (CAFs) in the tumor microenvironment was quantified by treating mono- and co-cultures with letrozole and then measuring the death rates.

**Results and Discussion:** Our simulations indicate that, even when a mutation is more drug resistant, its outgrowth can be delayed by abundant, low magnitude microenvironmental resistance across large regions of a tumor. These observations hold for different modes of microenvironmental resistance, including juxtacrine signaling, soluble secreted factors, and remodeled ECM. This result helps to explain the remarkable diversity of resistance mechanisms observed in solid tumors, which subverts the presumption that the failure mode that causes the quantitatively fastest growth in the presence of drug should occur most often in the clinic.

**Conclusion:** Our model results demonstrate that spatial effects can interact with low magnitude of resistance microenvironmental effects to successfully compete against genetic resistance that is orders of magnitude larger. Clinical outcomes of solid tumors are intrinsically connected to their spatial structure, and the tractability of spatial agent-based models like the ones presented here enable us to understand this relationship more completely.

## INTRODUCTION

Historically, drug discovery has been a largely empirical and predominantly serendipitous enterprise. However, modern targeted cancer therapies are rationally designed by integrating information on protein structure^46,58^, computational models of ligand binding^6,21^, and quantitative measurements in cell line models^40^. Because this approach is model driven and often rational, drug design is approached in a manner that is like classic engineering design. As a result of this rational design process, current drugs - approved in diseases that range from non-small-cell lung cancer (NSCLC)^14^ and breast cancer^22^ to chronic myelogenous leukemia (CML)^52^ - have been carefully crafted to avoid off-target toxicities and maximize therapeutic efficacy^13,19^.

A paradigmatic example of this therapeutic design cycle can be observed in the case of the first approved inhibitor (crizotinib) of the Anaplastic Lymphoma Kinase (ALK) fusions found in a subset of NSCLCs. Crizotinib was rationally repurposed from drug discovery efforts for a different kinase (c-Met)^10^. Following the clinical development of crizotinib, diverse point mutations in the ALK kinase were identified as the cause of resistance in a subset of patients^28,54^. Moreover, patients with brain metastases at baseline were found to have shorter progression free survival^9^. Thus, acquired drug resistance and blood-brain barrier penetrance limited the clinical effectiveness of first-generation therapy. These clinical findings emphasized the need for a second design cycle to create new drugs that have activity against resistance mutations and can cross the blood-brain barrier. These molecules (alectinib, brigatinib, and lorlatinib) have now been developed and approved, and they have subsequently increased progression free survival from ∼9 months to more than 33 months^27,55^. Thus, modern clinical cancer medicine uses cycles of design, failure-mode analysis, and testing to optimize cancer therapeutics for patients.

Because of these cycles of rational design and clinical testing, we view modern targeted therapies like other engineers view technologies like airplanes and bridges, i.e. as modern tools that exhibit a diversity of failure modes. While the biophysical design of the molecules that can target drug resistance is well rooted in first principles biochemical and biophysical concepts (i.e. the thermodynamics of ligand binding), the thermodynamics cannot quantitatively predict the output of biological evolution in patient tumors^35^. The diversity of evolutionary failure modes that are found in patients is lacking in explanations from current first principles evolutionary models. This is because classic results in theoretical cancer biology appear to imply that the diversity in the modes of clinical drug resistance should be smaller than what is observed, and that known microenvironmental/developmental resistance mechanisms should not be able to compete with genetic resistance^4,8,25,32^. Thus, a model-driven understanding of how the biological dynamics of intra-tumoral heterogeneity leads to clinical diversity is critically needed.

In this work we aimed to understand how different mechanisms of drug resistance with dramatically different scales of measured effects on cells can effectively compete in a single tumor, and thus eventually cause heterogeneous outcomes across a population of tumors. To do this, we parameterized a three-dimensional agent-based model of tumor growth. By varying the model rules to simulate different amounts of spatial constraint, we were able to explore parameter spaces that have predicted differences in the outcome of drug resistance evolution.

## RESULTS

### Clinical analyses reveal diverse evolutionary outcomes of drug resistance

We began by compiling counts of clinical resistance mechanisms for several targeted anticancer therapies and cancer types. The goal here was to get a tumor agnostic view of the range in interpatient diversity following treatment with targeted therapies. In binning cases by their mode of resistance (e.g., on-target single-nucleotide variants, SNVs), we identified a distribution of clinically significant resistance mechanisms within and across multiple cancers (Figure 1a).

**Figure 1:**
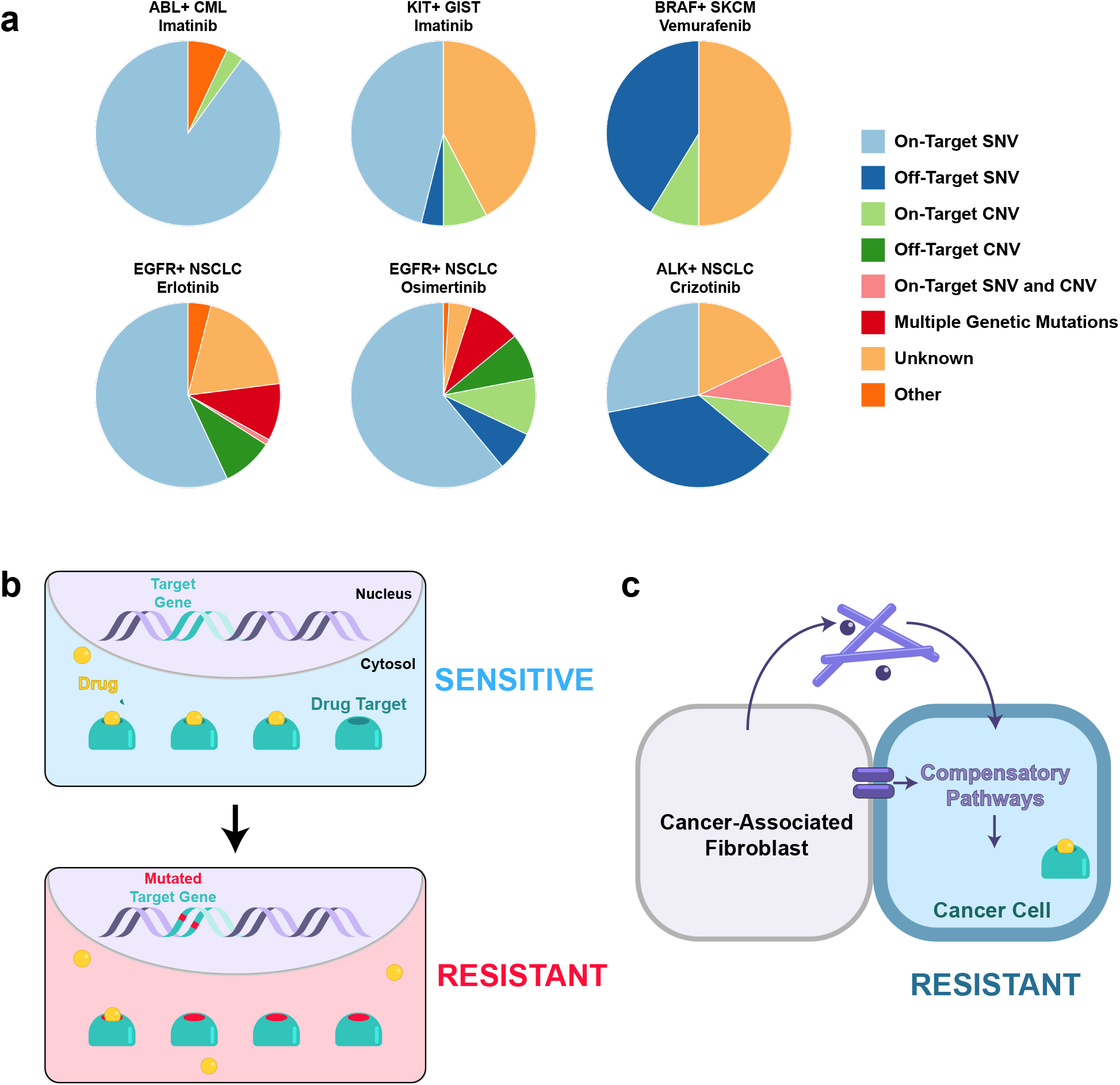
Mechanisms of anticancer drug resistance are diverse. (**a**) Pie charts indicating the distribution of resistance mechanisms for various drug-disease pairs. CML: chronic myelogenous leukemia; GIST: gastrointestinal stromal tumor; SKCM: skin cutaneous melanoma; NSCLC: non-small cell lung cancer. (**b**) Diagram of on-target genetic point mutations that interfere with drug binding and cause resistance. (**c**) Diagram of the multi-faceted roles CAFs can play in engendering phenotypic resistance in cancer cells.

These distributions indicate that resistance is often caused by genetic mutations. Given that targeted therapies are rationally designed to inhibit a target protein, point mutations that abrogate drug binding (on-target SNVs; Figure 1b) or amplification of the target gene (on-target CNVs) are commonly observed genetic events that create drug resistance^11,12,26,43,61^. Alternatively, cancer cells can also acquire genetic alterations outside the drug target (off-target SNVs and CNVs). These mutations often activate parallel pathways that compensate for pharmacologically-blocked oncogenic signals^63^.

However, in many patients, the mechanism of resistance is not identified by sequencing. These cases comprise the “unknown” category of drug resistance in our distributions. While such tumors may represent genetic forms of resistance that have not yet been elucidated, and thus are missed by genetic analyses, they can also arise from nongenetic modes of resistance, including epigenetic changes^62^ or an altered developmental state^2,29^. It is now well-established that these nongenetic factors can also be extrinsic to cancer cells, being conferred by the microenvironment^47,53,57^. Many noncancerous cells occupy the tumor microenvironment and can contribute to a drug resistant phenotype, including endothelial^17,44^ and immune cells^48,65^. However, among the most abundant stromal cells in tumors are cancer-associated fibroblasts (CAFs). These CAFs can alter tumor cell phenotypes in myriad ways^34,36,37^. During drug treatment, CAFs can promote resistance through juxtacrine signals^3,15^, secretion of soluble factors^49,59^, and remodeling of the extracellular matrix (ECM)^23,24^ (Figure 1c).

The suggestion of CAF-mediated resistance in the clinic, and the diversity of resistance mechanisms across tumors, is perhaps unsurprising on the surface. It is occasionally brushed aside in scientific discussions by arguing that intra-tumoral diversity should create inter-tumoral diversity. Clinically detectable tumors are large and heterogeneous, and it is hypothesized that at the initiation of treatment all possible single-nucleotide variants are already present in a clinically detectable tumor^4,7,31^. However, the observation that diversity within tumors portends diversity in outcomes across tumors belies basic evolutionary theory on clonal interference. Classic evolutionary dynamics results dictate that the fittest variants in a heterogeneous population should (on a long enough timescale) outcompete less-fit variants when a strong selective pressure like a cancer drug is applied. This leads to the natural question: if virtually all possible resistance variants exist in a tumor at the beginning of treatment, why does the single most resistant population not dominate relapse in each of these real-world parallel evolution experiments?

This paradox is compounded by the fact that the degree of fitness during treatment can vary widely across different modes of drug resistance. For example, preclinical evidence suggests that on-target SNVs can confer near-absolute resistance to a drug (>1000-fold increase in dose response curve inflection point over the wild-type drug target)^30^. In contrast, *in vitro* models reveal that cancer cells co-cultured with CAFs are granted only modest levels of drug resistance (1.5-to 3-fold increase in dose response inflection point)^64^. So how can low magnitude microenvironmental resistance ever outcompete high magnitude genetic resistance to drive relapse in patients? To address this, we first formalized back of the envelope calculations on tumor diversity and time to outgrowth into a well-mixed model of competition between microenvironmental resistance and genetic resistance *in vitro*.

### A well-mixed model of tumor evolution fails to explain clinical diversity and the clinical prevalence of low magnitude, microenvironmentally driven resistance

Relative subpopulation prevalence may provide an explanation for the ability of microenvironmental resistance to effectively compete with more fit, genetic resistance. If a sufficient number of tumor cells are endowed with resistance from the microenvironment upon treatment initiation, then even weak positive selective pressure may allow these abundant cells to dominate relapse before rarer genetic resistance has an opportunity to become detectable^35^. This hypothesis is bolstered by the fact that CAFs are often the most abundant stromal cells in the tumor microenvironment^56^, suggesting that many otherwise-sensitive cells could be endowed with a resistant phenotype via infiltrating CAFs.

To consider this possibility, we developed a well-mixed, stochastic model of tumor evolution to identify parameter regimes where CAF-mediated resistance can reach clinical detection before genetic resistance. Our birth-death-mutation model was parameterized using *in vitro* measurements of co-cultured cancer cells and CAFs (see Methods). In these simulations, an initially sensitive population of cancer cells expands until it reaches clinical detection (10^10^ cells), at which point drug treatment is initiated (Figure 2a). The division of sensitive cells can spawn genetically resistant subclones, which are unaffected by drug. Additionally, the expanding tumor actively recruits/activates CAFs. These CAFs support a subpopulation of otherwise-sensitive cells, and grant them low-level resistance during treatment^33^.

**Figure 2:**
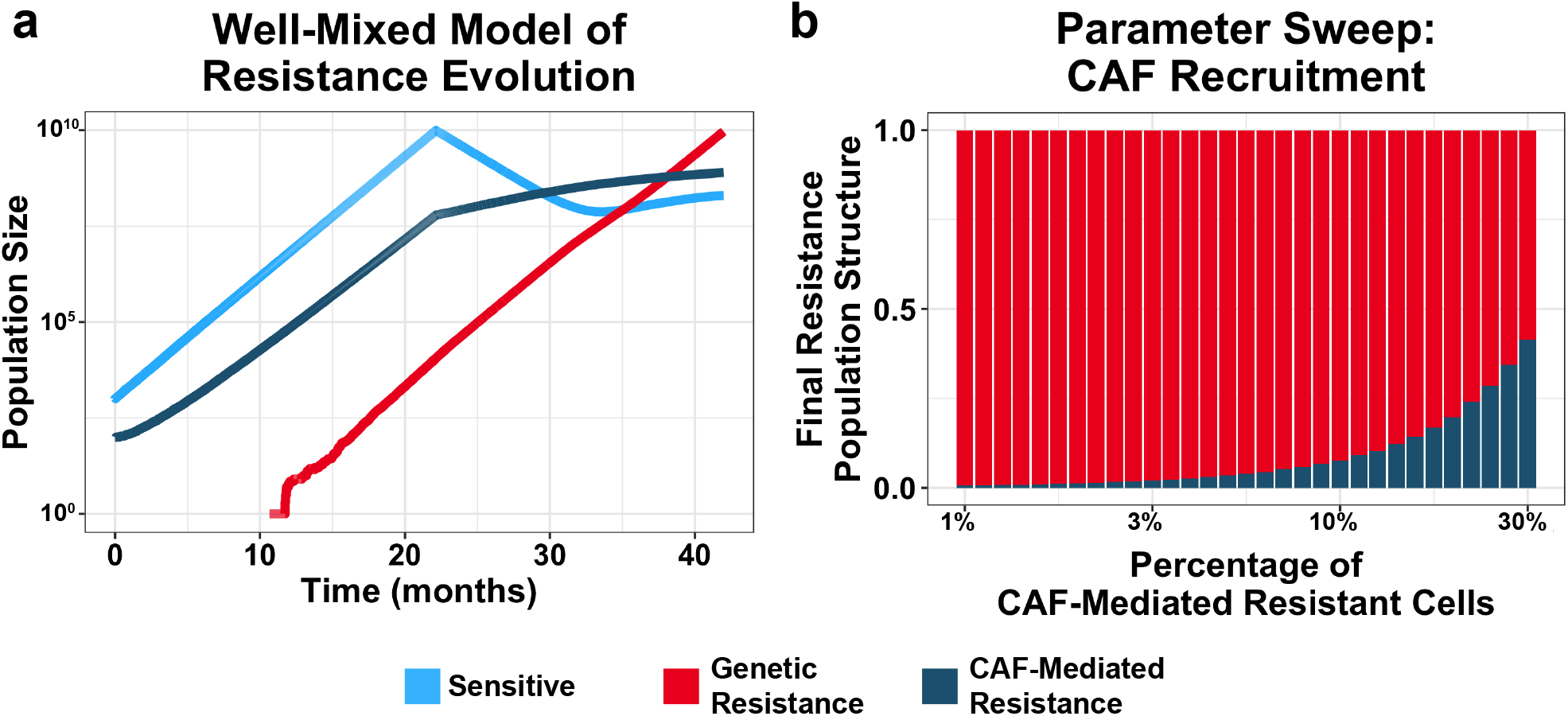
A well-mixed model of tumor evolution fails to explain the clinical observation of microenvironmental resistance. (**a**) Sample trajectory of tumor evolution in a stochastic, well-mixed model. A sensitive population (light blue) expands until clinical detection (10^10^), at which point drug treatment is initiated. Sensitive cells may spawn genetic resistance (red), and tumor cells recruit CAFs that support a CAF-mediated resistant population (dark blue). (**b**) Recruitment parameter sweep of the well-mixed model. Vertical bars represent the proportion of genetic and microenvironmental resistance upon relapse (10^10^ cells) for a range of recruitment parameter values. Proportions are the mean values for 48 simulations per condition.

By varying the rate of CAF recruitment, we can model how prevalent the CAF-supported populations must be at the beginning of treatment to reach clinical detection before genetic resistance. The simulation outcomes indicate that only for improbably high CAF recruitment rates can differences in abundances compensate for differences in fitness to favor low-level microenvironmental resistance over genetic resistance (Figure 2b). The results of this well-mixed model suggest that the observed prevalence of CAFs and the amount of resistance they confer is insufficient to explain its ability to outcompete genetic resistance in the clinic.

### An agent-based model enables tunable spatial competition

Prior work^35^ has demonstrated that restricted effective population sizes can alter the evolutionary landscape to favor earlier established (and therefore more abundant) populations over later-arising, more fit variants. This phenomenon explains the clinical distribution of resistance mutations in CML, where the effective population size is limited by the relatively small fraction of leukemic stem cells. Similarly, in adaptive therapy (the evolutionary-guided principle that reduced drug doses can extend time to disease progression), it has been shown that spatial competition in tumors can dramatically affect outgrowth dynamics^18,60^. We reasoned that, in the case of solid tumors, spatial competition could similarly act to limit the effective population size. In solid tumors, cancer cell division is limited by competition for space, nutrients and resources, and waste clearance. It is well-established that mitotic activity in solid tumors is limited to the tumor periphery, while the core is largely quiescent^18,20,42^. Thus, we hypothesized that these spatial determinants of competition could restrict a tumor’s effective population size and alter the evolutionary trajectories of microenvironmental and genetic resistance. Spatial factors may exaggerate the advantage of a more prevalent, but low-level resistance population over rarer, more fit subclones by simply limiting the space that a resistant cell has to divide. This hypothesis expands upon the theory of adaptive therapy, where moderate drug dosing maintains sensitive cells at steady-state levels, enabling competition to suppress the outgrowth of genetic resistance variants^1^.

To test this hypothesis, we developed a spatial agent-based model of tumor evolution to recapitulate spatial factors of competition. In our model, cells are randomly seeded on a 3D lattice and follow a stochastic birth-death process. To limit mitotic activity to the periphery, cell division in our model is limited by spatial availability. A dividing cell can search for an unoccupied site up to a certain distance, termed the *maximum budging distance*. If no available site is found, the cell does not divide, and the stochastic event is “skipped” (Figure 3a). However, if an unoccupied site is available, the dividing cell pushes nearby cells to occupy the free site in order to make room for the new daughter cell (Figure 3b). Thus, our model includes a spatial aspect to division; by decreasing the maximum budging distance, we can tunably limit mitotic activity to the tumor periphery and approximate the spatial competition present in a tumor (Figure 3c-f).

**Figure 3:**
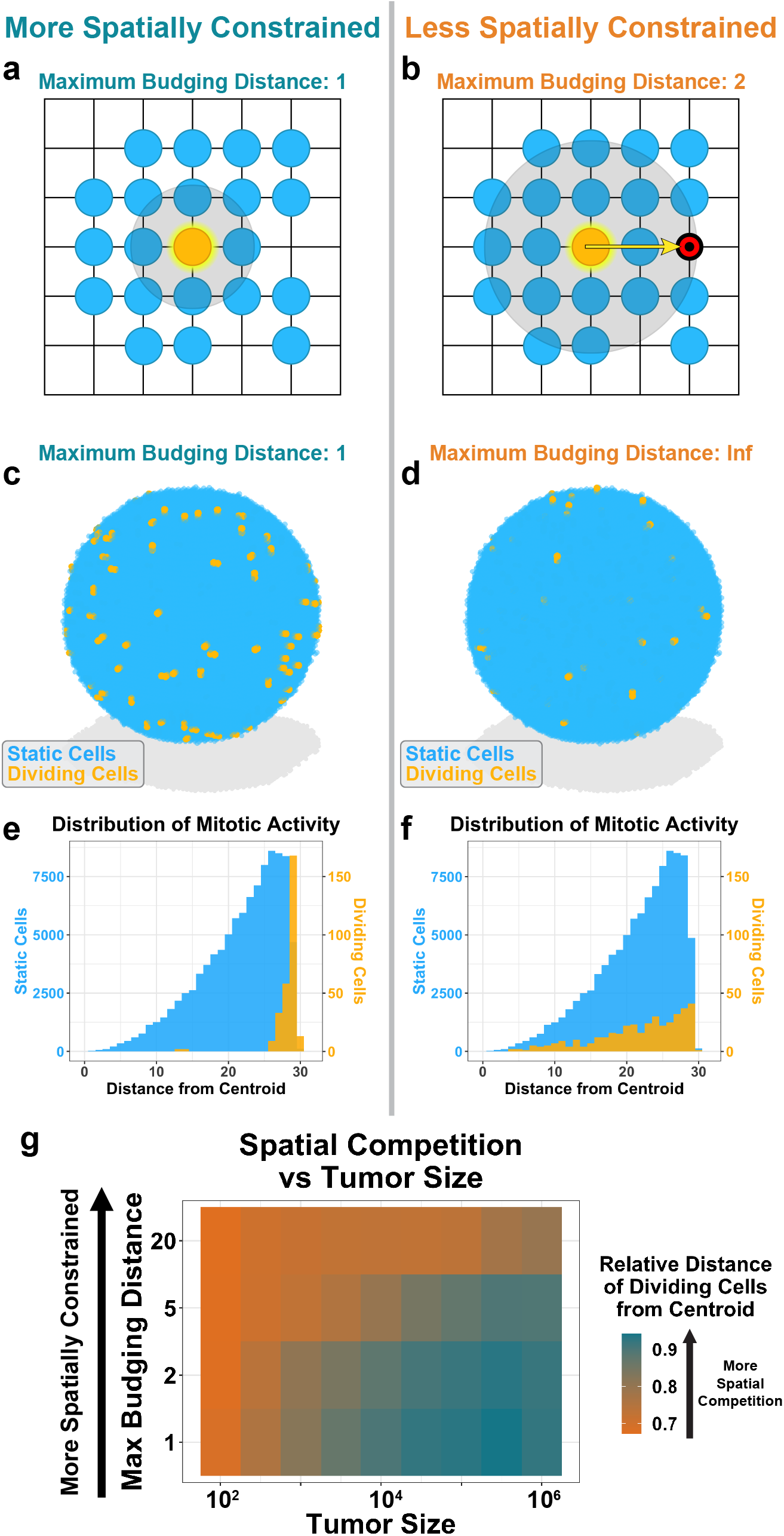
A spatial agent-based model enables tunable spatial competition. (**a**) Schematic of spatial agent-based model for a maximum budging distance of one. A cell is randomly selected to divide (orange). It searches for a region up to one unit length for unoccupied sites. (**b**) Spatial ABM schematic for a maximum budging distance of two. In this scenario, the dividing cell identifies an unoccupied site two units away. The algorithm draws a vector between the dividing cell and unoccupied site. The algorithm then budges cells along this vector to make room for the newly dividing cell. (**c**) Sample tumor simulation for a maximum budging distance of one. The simulation is run for 100 division events. Dividing cells are shown in orange. (**d**) Sample tumor simulation as in (**c**) for an infinite maximum budging distance. (**e**) Geographic distribution of dividing (orange) and static, nondividing (blue) cells for simulation shown in (**c**). (**f**) Geographic distribution of dividing (orange) and static (blue) cells for simulation shown in (**d**). (**g**) Spatial competition as a function of tumor size. Virtual tumors were seeded for a range of initial tumor sizes and maximum budging distances. Tumors were simulated for 100 division events. The average distance of dividing cells from the centroid (relative to the maximum distance from the centroid) was recorded. Values shown represent the average results for 48 instances. Orange indicates less spatial constraint; dark blue indicates more spatial constraint.

For the sake of computational efficiency, we used smaller tumor sizes (10^4^ cells) to represent larger, clinically detectable tumors. This approach is conservative, as spatial competition increases with tumor size (Figure 3g). Therefore, for equal proportions of resistance, tumors larger than those we simulate here will exhibit even greater degrees of spatial competition.

### CAF-mediated resistance and spatial competition can cooperate to suppress the outgrowth of genetic resistance

Having developed this spatial agent-based model, we revisited the question of competition between microenvironmental and genetic resistance. Alongside sensitive cancer cells, we introduced genetically resistant cancer cells and CAFs. In our model, CAFs were able to confer resistance to adjacent cancer cells. By varying CAF abundance and the maximum budging distance, we measured how microenvironmental resistance and spatial competition affect the evolutionary outcomes of these virtual tumors under treatment.

Our simulation results demonstrate that in the absence of CAFs, genetic resistance rapidly grows out and is the dominant population after one year (Figure 4a). With the introduction of CAFs, genetic resistance continues to comprise the majority of the relapsed tumor when spatial competition is low (i.e., for high maximum budging distances). These observations recapitulate the results of our well-mixed model. However, as spatial competition increases (i.e., for low budging distances), we find that low magnitude CAF-mediated resistance can effectively suppress the outgrowth of genetic resistance. For tumors with high degrees of CAF infiltration and strong spatial competition, these two factors can cooperate to maintain genetic resistance at very low abundances in the final population (Figure 4a) while the CAF-mediated resistant populations grow out (Figure 4b). These results may help to explain why microenvironmental resistance can come to dominate a real-world tumor in some patients, even when tumor size and mutation rate estimates suggest that potent genetic resistance is virtually guaranteed to exist at the time of clinical detection.

**Figure 4:**
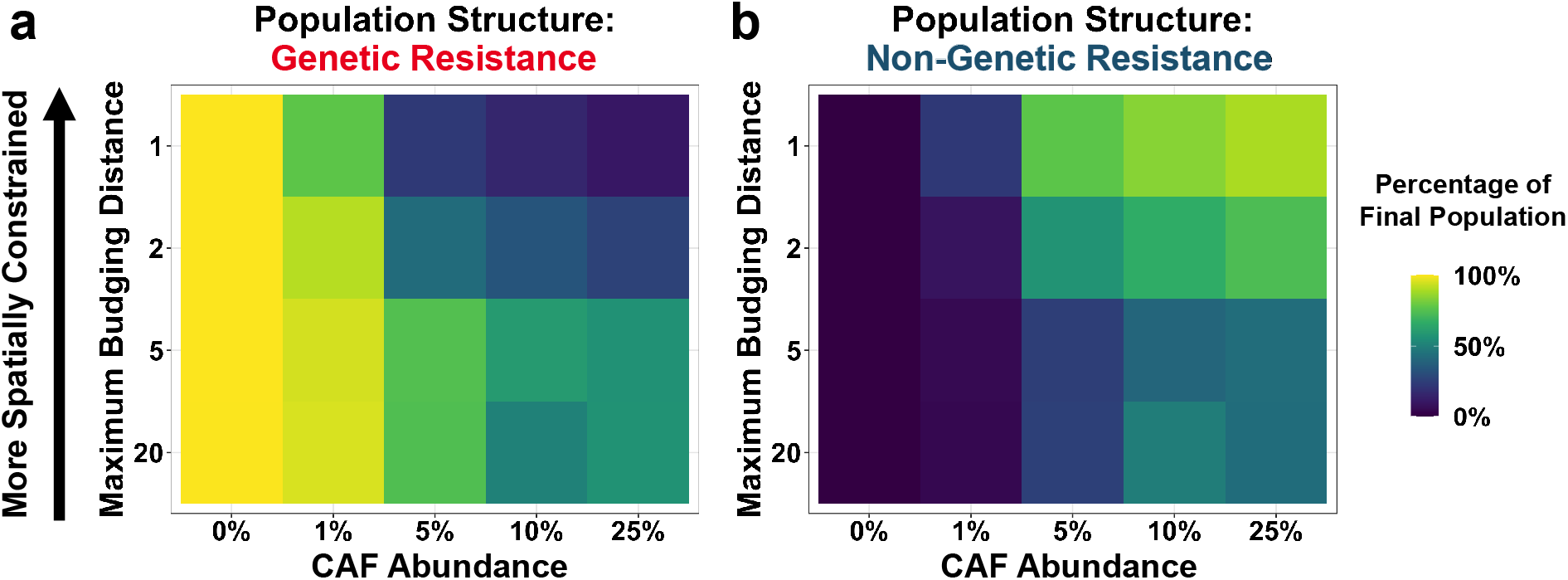
Spatial competition can limit the effective population size to favor microenvironmental resistance. (**a-b**) Outcome of CAF model simulations for varying CAF infiltration rates and maximum budging distances. The final population structure at 365 days was recorded. Values for each parameter combination represent the mean of 48 simulations. The fraction of cancer cells that are genetically resistant after one year are shown in (**a**). The fraction of cancer cells that are microenvironmentally resistant after one year are shown in (**b**).

### Paracrine signaling by CAFs can exaggerate the effects of spatial competition

Juxtacrine signaling of CAFs is a conservative base case that approximates a membrane bound cue. Beyond juxtacrine signaling, CAFs can secrete soluble factors that exert longer-range effects. These paracrine signals can modify the phenotypes of cancer cells at a longer distance and confer low magnitude resistance to cancer cells beyond those immediately adjacent to the CAF. We reasoned that these far-acting effects could amplify the ability of CAF-mediated resistance to suppress the outgrowth of genetic resistance in spatially competitive environments.

We modified our agent-based model to allow the influence of CAFs to extend beyond just immediately adjacent cells, by introducing a new parameter: CAF effect radius. Here we could study the effect of paracrine signaling on evolutionary outcomes (Figure 5a). Instead of modeling transport explicitly, we took a literature-based approach to estimating the largest radius that makes biological sense. Previous empirical studies have demonstrated that CAFs can affect cancer cells up to ∼100 microns away^38^, so we allowed the CAF effect radius to increase up to three cell lengths.

**Figure 5:**
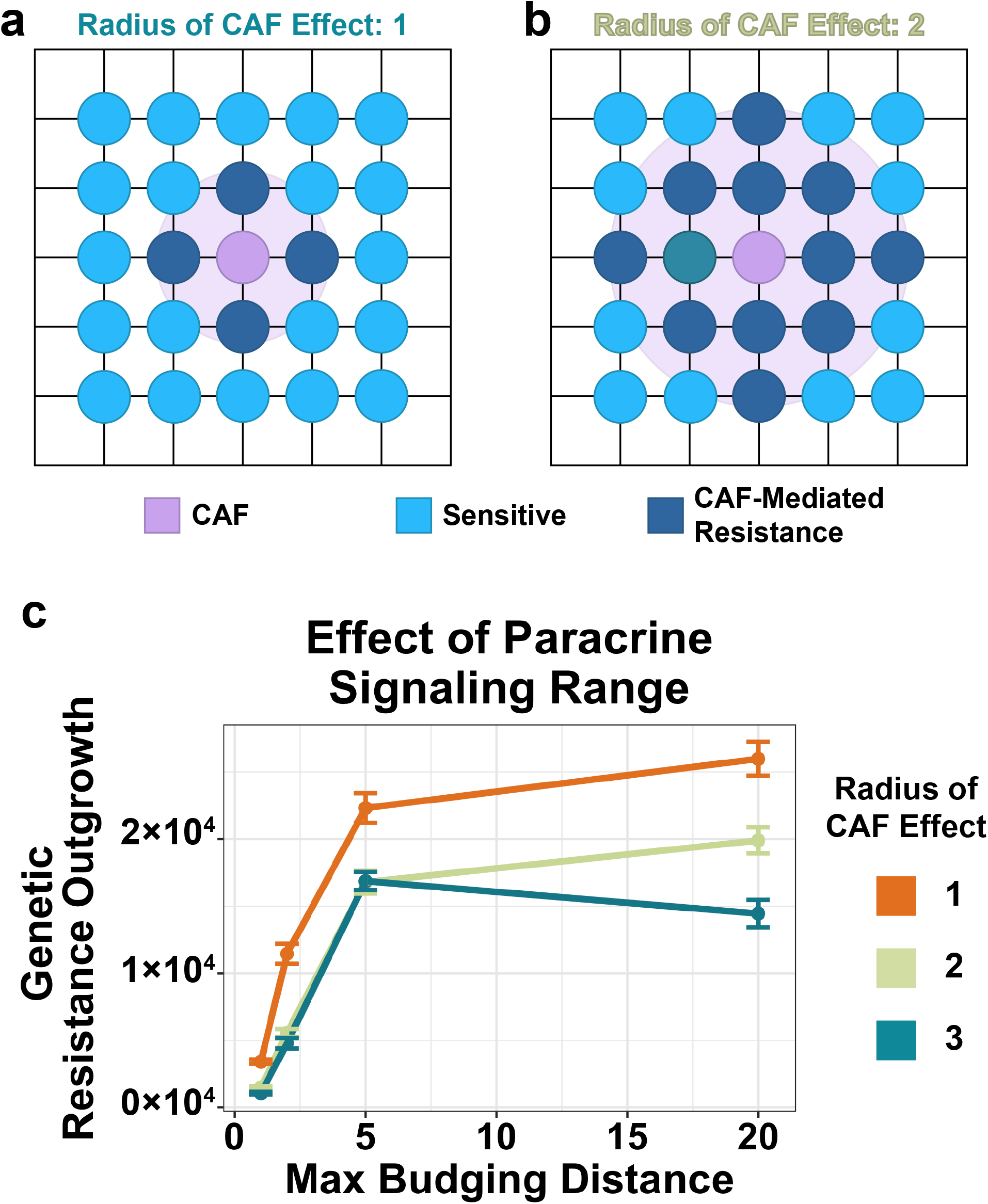
Longer-range paracrine signaling can amplify the effects of spatial competition on rare genetic resistance subclones. (**a**) Schematic of spatial agent-based model for paracrine signaling with a CAF effect radius of one. In this case, CAFs can confer resistance to cells up to one unit away. (**b**) Schematic for a CAF effect radius of two. In this case, CAFs can confer resistance to cells up to two units away. (**c**) Simulation results for the paracrine signaling model. The number of genetic resistance cells at one year are shown for a range of maximum budging distances and CAF effect radii. Points represent the mean of 48 simulations; error bars are standard error of the mean.

The simulation results indicate that as the CAF effect radius increases, the suppression of genetic resistance is amplified (Figure 5b). This trend was strongest for less spatially competitive environments, suggesting that longer-range paracrine signaling can compensate for weak competition to suppress the outgrowth of genetic resistance.

### Microenvironmental resistance conferred by remodeled ECM can restrict the outgrowth of genetic resistance

In addition to juxtacrine and paracrine signals, CAFs can also leave long-lasting effects on the tumor microenvironment by modifying the ECM. Migrating CAFs can deposit ECM components and secrete ECM-modifying enzymes, and these changes in external cues can help cancer cells escape therapeutic killing^23,24^. This mechanism of CAF-mediated resistance has the potential for even longer-range effects on cancer cells, as migrating CAFs leave a “wake” of resistance generating modified ECM.

Therefore, we hypothesized that ECM modifications could further augment the effects of spatial competition on favoring CAF-mediated resistance over genetic resistance. We modified our agent-based model to allow CAFs to actively migrate throughout the tumor. As CAFs move, they introduce modifications to the ECM; otherwise-sensitive cancer cells that are in contact with this modified ECM are granted low-level resistance (Figure 6a).

**Figure 6:**
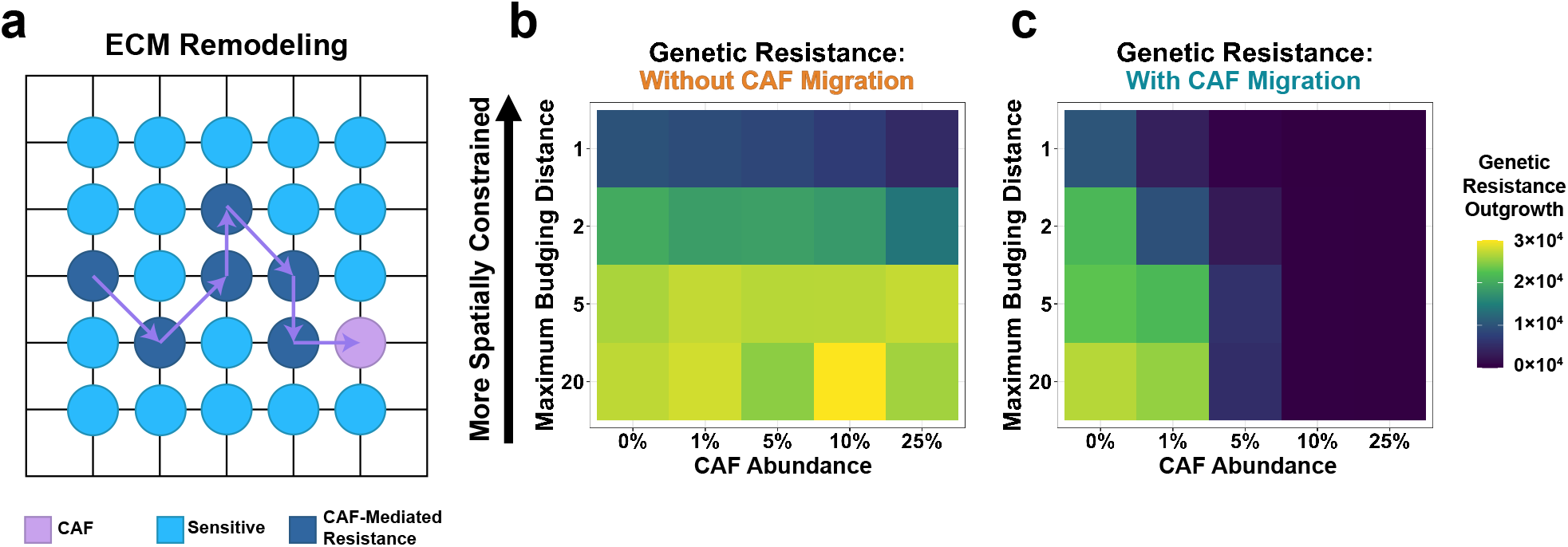
ECM-mediated resistance exaggerates the competitive suppression of genetic resistance. (**a**) Schematic of a migrating CAF remodeling of the ECM. In this model, CAFs move throughout the tumor, leaving a “wake” of modified ECM that confers microenvironmental resistance to otherwise-sensitive tumor cells. (**b-c**) Simulation results for the ECM model. The model was run for a range of initial CAF abundances and maximum budging distances. Values represent the mean genetic resistance population size at one year for 48 simulations. The model was evaluated for a migration rate of 0 (**b**) and 0.5 units/hr (**c**).

The simulation results indicate that in the absence of migration, ECM remodeling has very little effect on the final population structure of these virtual tumors (Figure 6b). However, migrating CAFs that modify the ECM can considerably shift evolutionary outcomes. ECM remodeling can exert long-range effects that severely restrict the outgrowth of genetic resistance; this is even true for tumors with moderately prevalent CAF populations (Figure 6c). In essence, the ECM remodeling and CAF migration magnifies the effect of the microenvironment in tumors with small CAF populations (1-5%).

In all scenarios explored here (juxtacrine signaling, paracrine signaling, and ECM remodeling), spatial competition can change the evolutionary competition to favor more abundant, low-magnitude microenvironmental resistance over rarer, but more potent genetic resistance. These theoretical results can help to explain the diversity of clinical resistance outcomes, suggesting that low-level resistance populations can exploit competition to suppress genetic resistance and drive treatment failure, even when genetic resistance is present at baseline.

## METHODS

### *In vitro* co-culture and model parameterization

BT20 cells and a panel of 16 CAFs derived from various tissues were cultured as described previously^33^. The *in vitro* net growth rate of BT20 cells were measured in the absence of CAFs. The average measurement across replicates gave a drug-free net growth rate of 0.0238 /hr. Co-culturing BT20 cells with CAFs was found to have little to no effect on their growth rates in the absence of drugs. The tonic death rate of BT20 cells was modeled using lag exponential death^16^ and estimated to be 0.001 /hr, suggesting a pure division rate of 0.0248 /hr. These values were used for the drug-free death and birth rates, respectively, of cancer cells in our model.

Next, the death rate of BT20 cells was measured in the presence of 100 nM letrozole. This measurement informed the drug kill rate for sensitive cells in our model: 0.0410 /hr (a net growth rate of -0.0172 /br). Death rate measurements were repeated in the presence of one of 16 CAFs sourced from various tissues^33^. The mean co-cultured death rate measurements informed the drug kill rate for CAF-mediated resistant cells in our model: 0.0165 /hr (a net growth rate of 0.0073 /hr).

### Description of well-mixed model

We developed a stochastic model of tumor evolution with well-mixed subpopulations that follow a birth-death-mutation process. The model considered an initial population of 10^3^ sensitive cancer cells (***S***), which expand until clinical detection (10^10^ cells), when treatment is initiated. Dividing cancer cells may spawn genetically resistant cells (***R***) with a frequency of 10^−7^ /division. In addition, cancer cells actively recruit CAFs to the tumor. CAFs support a fraction of the sensitive cell population termed adjacent cells (***A***), granting resistance that allows them to maintain a positive net growth rate during treatment. This fraction of otherwise-sensitive cells supported by CAFs is a function of the maximum number of adjacent cells, *A*_*max*_. This quantity is in turn a function of the product of the CAF recruitment rate and the number of cancer cells one CAF can support, a term collectively referred to as the *recruitment parameter* in the model. While adjacent cells divide exponentially in the model, only a proportion of daughter cells (1 − ***A***/*Amax*) are assigned to the adjacent class. The remainder of daughter cells, spawned at a rate of

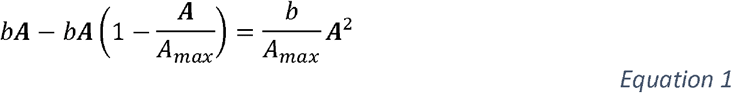

(where *b* is the division rate), are instead designated as sensitive.

This system was solved stochastically using the Gillespie algorithm with adaptive tau leaping^5^ in MATLAB. The final propensity vector and state change matrix was

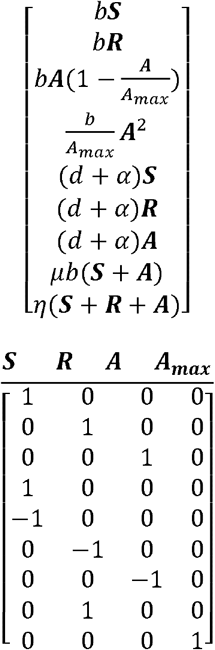

where, ***S, R***, and ***A***, are the sensitive, resistant, and adjacent populations, respectively; and *b, d, α, μ*, and *η* are the division, death, phenotype-specific drug kill rate, mutation rate, and recruitment parameter, respectively. The recruitment parameter was allowed to vary in order to determine the adjacent population size required at treatment initiation to effectively compete with genetic resistance.

### Description of base agent-based model

We developed a spatial agent-based model of competing tumor cell subpopulations. In our model, cells are randomly seeded on a three-dimensional lattice. Cells follow a birth-death process, and the model is solved exactly using a modified Gillespie algorithm. In a single time step of the simulation, if a division event occurs, a cell is randomly drawn from the appropriate population. The algorithm searches for unoccupied sites in the 3D lattice adjacent to the dividing cell. If an unoccupied site is defined, the cell divides and the new daughter cell occupies the previously vacant site. If no such site is identified, the algorithm expands its search around the dividing cell by one unit and searches for unoccupied sites in this region. This process continues until a vacant site is identified or the maximum budging distance is reached; if the maximum budging distance is reached this is an unproductive event and no cell division occurs. Thus, a small maximum budging distance limits cell division to the tumor’s periphery while maintaining a quiescent core.

If an unoccupied, non-adjacent site is identified before the maximum budging distance is reached, then the dividing cell pushes cells (i.e., the algorithm “budges” cells) to make room for the new daughter cell. In this scenario, the algorithm draws a vector between the dividing cell and the unoccupied site. It then identifies the cell adjacent to the unoccupied site that is nearest the vector (calculated using the parallelogram definition of the vector cross-product), excluding cells further from the dividing cell than the unoccupied site is. The algorithm then moves this identified cell to the unoccupied site, generating a new unoccupied site where it had been. This process repeats until the vacant site is adjacent to the dividing cell, at which point the division event occurs and the newly spawned daughter cell occupies the vacant site.

To measure the effects of the maximum budging distance and initial population size on spatial competition in the model, these parameters were allowed to vary. For each parameter combination, 48 tumors were simulated for the first 100 division events. The geographic locations of these dividing cells were recorded for each simulation. Then, the ratio of their average distance from the tumor centroid was divided by the maximum distance from the tumor centroid, yielding a measure of spatial constraint.

### CAF juxtacrine model description

The base model described in the previous section was modified to include sensitive, resistant, and CAF-supported (i.e., adjacent) cancer cells, as well as CAFs. For the sake of computational efficiency, simulations began at the onset of treatment with 10^4^ randomly seeded cells. Otherwise-sensitive cancer cells immediately adjacent to CAFs were labeled adjacent. This designation is determined after each iteration of the simulation, to account for budging that may occur during cell division. The birth, death, and drug-kill rates were identical to those parameters used in the well-mixed model. The initial proportion of CAFs and maximum budging distance were allowed to vary. For every parameter combination, the simulation was run 48 times and the final population structure at 365 days was recorded.

### CAF paracrine model description

The juxtacrine model described in the previous section was modified to consider the effects of paracrine signaling. In this model, the CAF effect was not limited to immediately adjacent cancer cells. Rather, cells up to the distance of the *radius of CAF effect* parameter were labeled as CAF-supported and received the same modest drug resistance phenotype. The largest CAF effect radius simulated was three cell lengths, guided by *in vitro* measurements of longer-range CAF effects^38^. The radius of CAF effect and maximum budging distance parameters were allowed to vary, while the initial CAF proportion was held constant at 5%. For every parameter combination, the simulation was run 48 times and the final population structure at 365 days was recorded.

### ECM remodeling model description

The base spatial agent-base algorithm was modified to model the effects of ECM remodeling. In this model, CAFs were able to freely migrate throughout the tumor. When a CAF was selected to migrate, it would randomly select an adjacent site to occupy. The site previously occupied by the CAF was then labeled “remodeled ECM”, and any cancer cell that came to occupy that site would be designated as microenvironmentally resistant. The CAF migration rate was set to 0.5 cell lengths/hr, guided by literature estimates of CAF migration^39^. The ECM remodeling was transient, and sites would lose their resistance-conferring state at a rate of 0.05 /hr. The maximum budging distance and CAF infiltration rates were allowed to vary, and the final population structure at 365 days was recorded. Our code is available at https://github.com/pritchardlabatpsu/ABMResistanceEvolution.

## DISCUSSION

The scale and extent of intratumoral heterogeneity has been unlocked by a revolution in genomic technologies^50,66^. The diversity of tumor heterogeneity and the diversity of the clinical mechanisms of drug resistance is now well established^35,41^. Ultimately, the biological forces that govern intratumoral competition, when aggregated across the population, are responsible for interpatient heterogeneity, but these forces are poorly understood^45^. Previously, work on the failure modes of ABL kinase inhibitors^35^ has led to a simple principle: when evolution favors the most mutationally probable failure mode, then so should drug design. By identifying the biological mechanisms that govern the clinical prevalence of drug failure modes, we can identify optimization criteria for further drug design. But beyond single amino acid changes in drug targets, the diversity of resistance mechanisms in solid tumors, from genetic to micro-environmental mechanisms requires more analysis.

It is easy to qualitatively argue that because a tumor is diverse, then the mechanisms of resistance across patients should be diverse too. However, back of the envelope calculations reveal interesting treatment paradoxes that need deeper biological inquiry. In a large tumor with 10^8^-10^12^ cells and a ≥10^−8^ /division mutation rate, it seems that every possible resistance mutation should be present at the start of therapy^4,7,31^. Measurements of mutational diversity in real human tumors support these calculations^51^. Further coarse calculations suggest that when a drug kills 50-99% of tumor cells over 3-12 months of therapy, all clinically detectable tumors should inevitably lead to the outgrowth of a single high strength resistance mutant in every patient. Biologically, we know that resistance mutations that are this potent do exist, but clinically we know that the diversity of observed resistance mechanisms is extremely high. The reason for this discrepancy is understudied. Elucidating the possible mechanisms by which intratumoral heterogeneity can control interpatient heterogeneity is critical to understanding the nature of drug resistance and how to stop it with new drugs.

Previous research in adaptive therapy has identified spatial heterogeneity as a possible reason for the efficacy of adaptive therapy regimens^1,18,60^. Because of work in the adaptive therapy field, and our biological intuition, we suspected that spatial competition in the tumor bed might dramatically change clonal competition between rare and highly drug resistant mutations that confer complete drug resistance, and the low magnitude resistance induced by microenvironmental cues present across large fractions of a tumor. This clonal competition between rare and common is interesting because in the presence of spatial constraints, extremely poor drug resistance that is cell extrinsic and orders of magnitude weaker can effectively compete highly potent resistance mechanisms.

Interestingly, across all of the simulations in this paper, amplifying the spatial effect of a CAF population by moving from juxtacrine to paracrine signaling simulations (Figures 4 and 5), or by including migratory cells that can remodel ECM as they travel (Figure 6) amplifies the evolutionary effects of the relatively small growth advantages supplied by microenvironmental resistance. To our knowledge, this is the first study to show that small effect size resistance (1.5-3 fold changes in drug sensitivity) can in theory compete with large effect size resistance (1,000 fold changes in drug sensitivity) if strong spatial competition exists. This strong spatial competition is achieved in this study by controlling a budging parameter. Intuitively, this budging parameter is a surrogate for numerous biological and physical phenomena. Low budging distances might represent tumors that are less migratory, or stiffer, with a less degradable ECM. High budging distances may be modeling environments where tumor proteases are more capable of facilitating movement, or where tumor cells are more biologically migratory. While future work is necessary to identify which measurable parameters in human tumors correlate with the idea of a budging distance, identifying these markers in human tumors that provide reliable estimates of spatial competition that may be useful to identify which patients are likely to develop microenvironmental resistance in the clinic.

## Acknowledgements

JRP, ML, and SRP acknowledge support from the NCI U01CA265709-01. ML, SRP, and JRP acknowledge research support from the JKTG foundation for health research.

